# The smallest infectious substructure encoding the prion strain structural determinant revealed by spontaneous dissociation of misfolded prion protein assemblies

**DOI:** 10.1101/2023.03.21.533631

**Authors:** Jan Bohl, Mohammed Moudjou, Laetitia Herzog, Fabienne Reine, Fiona Sailler, Hannah Klute, Frederic Halgand, Guillaume Van der Rest, Yves Boulard, Vincent Béringue, Angelique Igel, Human Rezaei

**Author notes:** **Author contributions:** A.I., V.B. and H.R. designed the research; all authors contributed to the research and analyzed the data; and V.B., J.B., A.I. and H.R. wrote the paper. The authors declare no competing interests. **Declaration of Competing Interest:** The authors declare that they have no known competing financial interests or personal relationships that could have appeared to influence the work reported in this paper. Corresponding authors and senior authorships: Vincent Béringue, <u>Angélique Igel</u>, Human Rezaei, MAP2, VIM, INRAe, Domaine de Vilvert, 78352, Jouy-en-Josas, France.

## Abstract

It is commonly accepted that the prion replicative propensity and strain structural determinant (SSD) are encoded in the fold of PrP^Sc^ amyloid fibril assemblies. By exploring the quaternary structure dynamicity of several prion strains, we revealed that all mammalian prion assemblies exhibit the generic property of spontaneously generating two sets of discreet infectious tetrameric and dimeric species differing significantly by their specific infectivity. By using perturbation approaches such as dilution and ionic strength variation, we demonstrated that these two oligomeric species were highly dynamic and evolved differently in the presence of chaotropic agents. In general, our observations of seven different prion strains from three distinct species highlight the high dynamicity of PrP^Sc^ assemblies as a common and intrinsic property of mammalian prions. The existence of such small infectious PrP^Sc^ species harboring the SSD indicates that the prion infectivity and the SSD are not restricted only to the amyloid fold but can also be encoded in other alternative quaternary structures. Such diversity in the quaternary structure of prion assemblies tends to indicate that the structure of PrP^Sc^ can be divided into two independent folding domains: a domain encoding the strain structural determinant and a second domain whose fold determines the type of quaternary structure that could adopt PrP^Sc^ assemblies.

**Highlights:**

- Mammalian prion assemblies are highly dynamic
- Prion assemblies spontaneously disassemble into two infectious oligomers
- Prion infectivity is not exclusively encoded in the amyloid fibrils’ structure
- Two independent folding domains could structure Prion assemblies

## Introduction

The prion paradigm unifies the transmission of several age-related, incurable neurodegenerative disorders based on assisted catalytical protein refolding concerted with the acquisition of quaternary structures [1]. In principle, the prion paradigm consists of an autocatalytic structural switch of a host-encoded monomeric protein or peptide induced by the same protein or peptide in an aggregated conformation [2, 3]. This aggregated conformer or assembly plays the role of a template. During this assisted structural switch, the biological information encoded in the structure of the assemblies is transferred to the monomeric protein (templating step), leading to the perpetuation of the biological information [4]. In human and animal transmissible spongiform encephalopathies or prion diseases, the host-encoded cellular prion protein (PrP^C^) undergoes an induced structural rearrangement into a catalytical active, pathological conformation called PrP^Sc^, which serves as a template for de novo conversion of PrP^C^ into PrP^Sc^ in an autocatalytic process [5, 6].

For conventional infectious diseases where variations in the clinical manifestation of the disease define the pathogenic strain, in prion diseases, variations in the incubation period, neuropathological patterns and biochemical properties of PrP^Sc^ assemblies differentiate prion strains [7, 8]. At the molecular level, these biological differences are the consequence of differences in the structure of PrP^Sc^. How the strain structural determinant (SSD) is encoded in the fold of PrP^Sc^ assemblies remains unclear, despite the structure of protease-resistant PrP^Sc^ assemblies from four strains being currently known [9–12]. Nonetheless, the SSD governs the biochemical properties of PrP^Sc^ assemblies, such as the type of PrP^Sc^ fragments generated after proteolysis [2, 13], apparent resistance to unfolding [14, 15], and size distribution of PrP^Sc^ assemblies at the terminal stage of the disease [16–18].

In the prion literature, there is compelling evidence that different strains can be propagated on one given primary structure [19–21]. This indicates that multiple SSDs can be encoded in one PrP primary structure. This assumption is particularly true if we exclude the contribution of posttranslational modifications of the primary structure to the SSD, as evidenced by the similarity of the N-glycans attached to PrP^Sc^ from two distinct strains propagating on the same PrP primary structure [22]. Based on our own experimental transmission data, more than fifteen different prion strains can be stably and distinctively propagated on the sheep PrP sequence (V_136_R_154_Q_171_ allele) (Fig. 1 and SI.1), indicating that one given primary prion structure can adopt at least fifteen stable and different conformations. How such a broad diversity is structurally encoded at the scale of PrP^Sc^ assemblies remains undetermined and conceptually difficult to reconcile with the current models of prion assemblies. This question is particularly interesting when considering the large number of mammalian prion strains. At which scale –, i.e., primary, secondary, tertiary, quaternary or supraquaternary PrP^Sc^ assembly structures, such a broad spectrum of structural information that can be encoded remains elusive.

**Figure 1.**
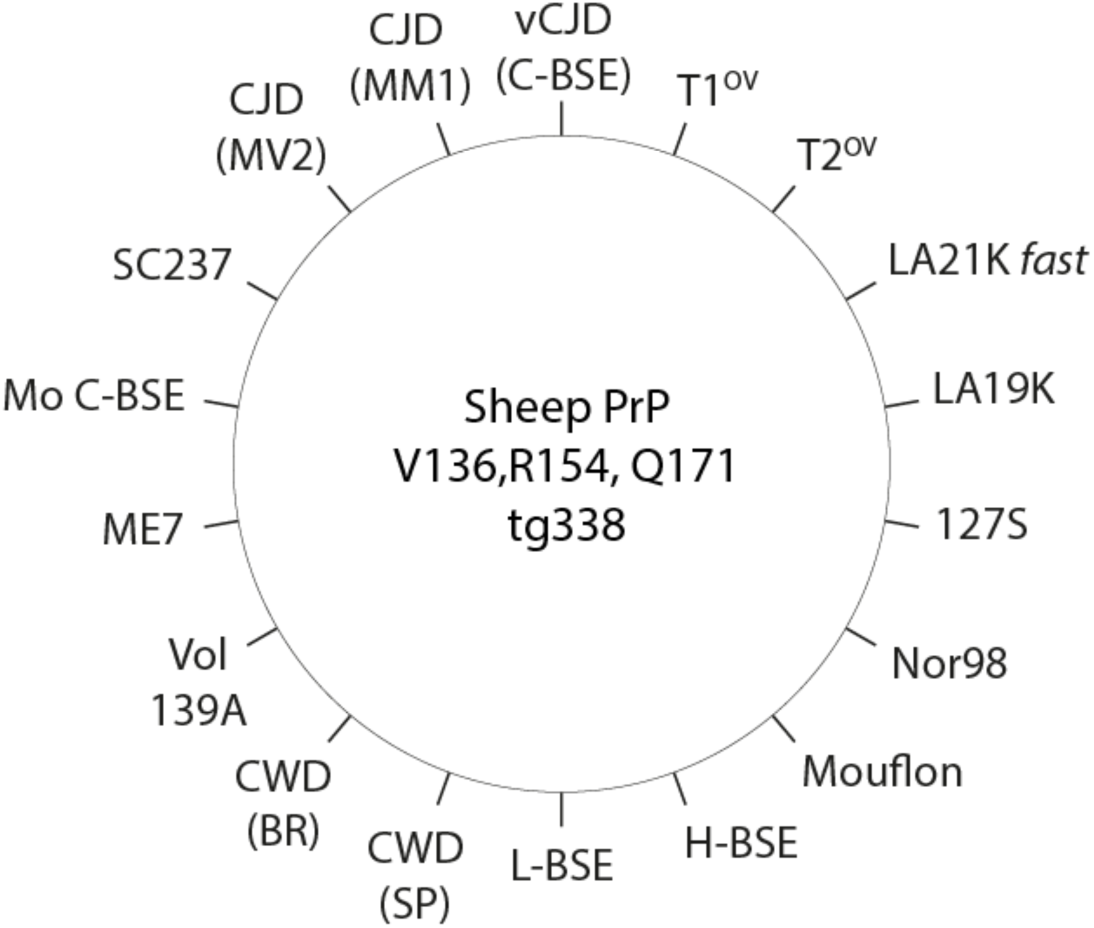
Different prion strains stabilized on tg338 mice expressing the allelic variant V136, R154, Q171 of sheep PrP^C^. As shown, sheep PrP^C^ can be refolded by serving as a substrate for more than 15 different prion strains. Each fold is at the origin of a given strain structural determinant (SSD) and at the origin of the strain-specific incubation time (see SI 1). The fact that a given primary structure of PrP^C^ (here sheep V136,R154,Q171 variant) is able to be refolded in different structures highlights the multistability of PrP.

Putting aside the strain dimension, multiple PrP^Sc^ conformations coexist within a given prion strain [23-25]. Even if the extent of this diversity is not yet well established, it refers to the quasispecies concept, which has been proposed to be at the base of prion adaptation and strain evolution [23]. The existence of multiple conformers within a strain and, more specifically, biologically cloned strain adds another level of structural complexity to PrP^Sc^ assemblies and questions whether intrastrain structural diversity involves the SSD or another PrP^Sc^ structural subdomain. Recent investigations demonstrated that, independent of the prion strain, deterministic structural diversification occurred during the replication process, leading to the formation of two sets of protease-resistant prion assemblies called PrP^ScA^ and PrP^ScB^ [26]. While for a given prion strain, PrP^ScA^ and PrP^ScB^ assemblies based on their specific infectivity are structurally different, they converge to the same strain. This suggests that different sets of assemblies from a given prion strain share the same SSD.

To determine at which PrP^Sc^ assembly structural scale the SSD is encoded and the potential variations among strains, we established a native method to disassemble PrP^Sc^ from seven different cloned prion strains into their smallest unit. We showed that, as a generic process, PrP^Sc^ from the seven prion strains disassembled into two small oligomer species, called P1 and P2. Using different size estimation methods, we determined the sizes of P1 and P2 to be tetramers and dimers, respectively. The comparison between the specific infectivity of P1 and P2 for three different strains revealed significant differences, suggesting the existence of at least two intrastrain structural subpopulations. The existence of two intrastrain subpopulations has been further confirmed by using chaotropic treatment, leading to the segregation of the initial PrP^Sc^ population into high molecular weight assemblies devoid of infectivity and an infectious small oligomeric assembly.

## Results

### Disaggregation of PrP^Sc^ assemblies into infectious small oligomeric assemblies

To determine how prion strain information is encoded in the PrP^Sc^ structure, we established native disassembling conditions to disaggregate PrP^Sc^ assemblies into their elementary subunit suPrP. Brain homogenates (BHs) of seven different prion strains from hamster and transgenic mice expressing ovine (tg338) and mouse (tga20) PrP were proteinase K (PK)-digested to remove PrP^C^ before a 4-hour solubilization step at 37 °C in a buffer containing dodecyl maltoside and sarkosyl at 5 mM and 50 mM, respectively (see M&M for more details). At these concentrations, these two detergents do not significantly affect either protein tertiary structure [27, 28] or prion infectivity [16, 18, 26]. The solubilization procedure led to a translucid solution. After a subsequent centrifugation step at 15 000 x *g,* the supernatants and the pellets were collected and analyzed for PK-resistant PrP^Sc^ (PrP^res^) content by western blotting. The pellets contained negligible amounts of PrP^res^ compared to the supernatants (see SI2). The supernatants were then directly analyzed by size exclusion chromatography (SEC) using a running buffer that did not contain any detergent to avoid the formation or maintenance of lipidic micellar structures that could interfere with the intrinsic elution volume of PrP assemblies [28, 29]. The collected fractions were analyzed by western blotting for PrP^res^ content. As typically shown for cloned ovine LA21K *fast*, LA19K, and hamster 263K prion strains, the chromatograms revealed the existence of two discreate and well-defined PrP^res^ peaks called P1 and P2, indicating the presence of at least two sets of oligomeric PrP^res^ assemblies (Fig. 2A-C). For LA21K *fast*, LA19K and 263K, the SEC analyses of mouse ME7, 139A, 22L, and Fukuoka-1 prion strains revealed similar elution profiles. Only the relative proportions of P1 and P2 varied among the strains analyzed. The seven strains tested here thus share a common disassembly process generating P1 and P2 (Fig. 2D). According to our calibration, the hydrodynamic radius of the assemblies in P1 and P2 ranged between a trimer and tetramer of PrP for P1 and a PrP dimer for P2. Each injection was repeated at least three times, with only minor deviations in the elution profiles (see error bars in Fig. 2), demonstrating the high repeatability of the SEC experiments.

**Figure 2:**
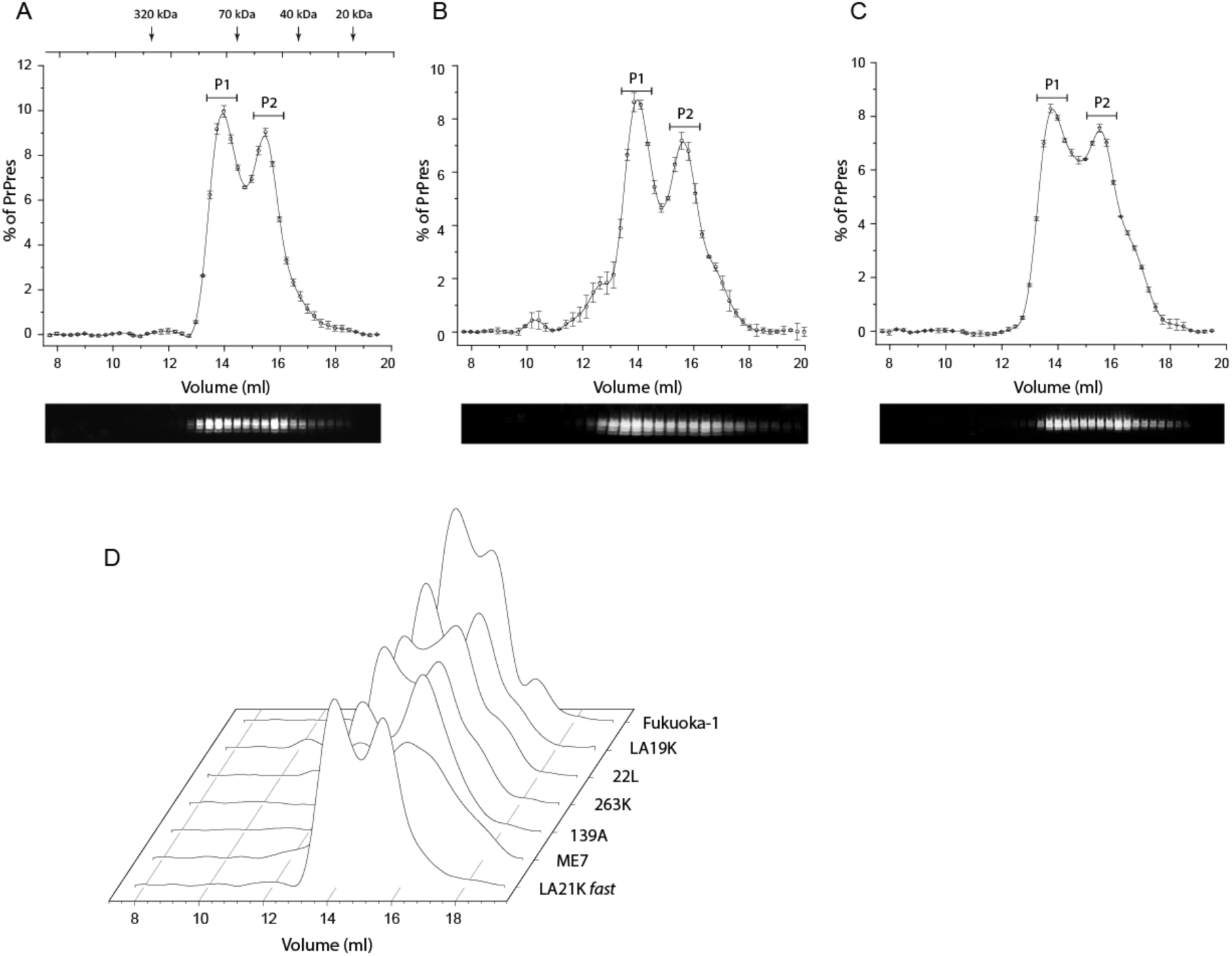
Analysis of the quaternary structure of PrP^res^ assemblies by SEC. Chromatograms of solubilized PK-treated brain homogenates containing LA21K *fast* (A), LA19K (B) and 263K (C) prions. The mean levels of PrP^res^ per fraction were obtained from the immunoblot analysis of *n*=3 independent SEC. Representative western blots used for PrP^res^ level quantification are presented below the chromatograms. For the three strains, the chromatograms revealed the presence of two discreet peaks called P1 and P2. The same analysis for four other strains (ME7, 139A, 22L and Fukuoka-1, (D).

To determine whether purified infectious fibrillar PrP^Sc^ follows a disassembly pathway similar to that of PrP^Sc^ present in the brain homogenate, we first purified 263K assemblies according to Wenborn and colleagues’ protocol [30]. The hydrodynamic radius as well as the mean average molecular weight (<MW>) of these purified assemblies as estimated by static (SLS) and dynamic light scattering (DLS) revealed a <MW> of 8 MDa with a hydrodynamic mean average radius of the assemblies centered at approximately 150 nm (Fig. 3A). The purified 263K at 20 nM (concentration expressed equivalent to monomer) was then incubated at 37 °C under identical conditions as for BH (Fig. 2C) before analysis by SEC coupled to SLS. As shown in Fig. 3A and B, when initially the size of purified 263K was approximately 150 nm with a <MW> of approximately 8 MDa, after solubilization, SEC revealed a single unique peak eluting at the P2 position, with a mean average molecular weight of ≈ 54 *kDa* (Fig. 3B), which is in good agreement with a dimeric N-terminal truncated mixture of mono- and biglycosylated PrP. This observation indicates that after solubilization, the very large assemblies initially present in the purified products were transformed into P2 species. However, when PrP^Sc^ present in the brain homogenate gave rise to two peaks, P1 and P2, the disassembly of purified 263K gave a unique peak eluting at the P2 position.

**Figure 3:**
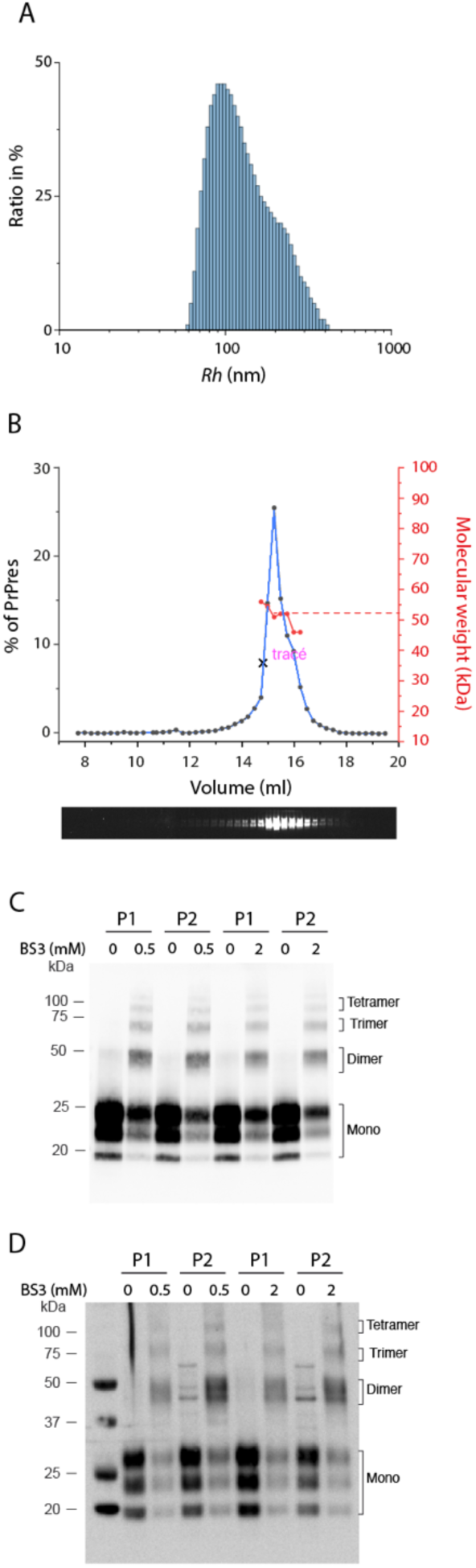
Size characterization of P1 and P2 assemblies. (A) Hydrodynamic radius (Rh) of purified 263K assemblies as estimated by static light scattering (SLS). (B) SEC analysis of purified 263K assemblies solubilized under the same conditions as for brain homogenate. The light scattered intensity measurement of the fractions corresponding to the P2 peak allows us to estimate the <MW> of assemblies eluting at the P2 position. By neglecting the edge effect, the maximum of the peak corresponds to <MW> of 54 kDa, compatible with a PrP^res^ dimer core. (C-D) western blot profile of SEC fractions corresponding to P1 and P2 from 263K (C) and LA21K *fast* (D) PrP^res^ before and after crosslinking with different concentrations of BS3 (0.5 and 2 mM). BS3 crosslinking led to the appearance of mainly a dimeric band even at high concentrations. Independent of BS3 concentration, other minor PrP multimeric bands, such as trimer and tetramer bands, were also observed, suggesting a dynamic exchange between higher quaternary structure arrangements [47].

To precisely determine the molecular weight of PrPres assemblies present within the P1 and P2 peaks, a crosslinking approach was adopted using BS3 as an 11 Å bifunctional amine crosslinker to covalently trap the oligomeric state. Collected fractions corresponding to P1 and P2 from 263K and LA21K *fast* were incubated with 0.5 or 2 mM BS3 to covalently crosslink lysine residues distant below 11 Å. The analysis of crosslinked P1 and P2 by western blot revealed the existence of dimeric bands (Fig. 3C and D) for both strains. Other minor PrP multimeric bands, such as trimer and tetramer bands, were also observed. However, their ratio did not evolve with higher amounts of BS3, indicating that the dimeric species in the P1 and P2 peaks are preferentially crosslinked by BS3.

Thus, according to the hydrodynamic radius estimation, the mean average molecular weight determination by SLS and the crosslinking experiments, the quaternary structure of P2 species corresponds to a dimer. According to their hydrodynamic radius, the size of objects forming the P1 peak are expected to range between a trimer and a tetramer of PrP. However, the crosslinking experiment tends to indicate a dimer. This discrepancy can be explained if P1 corresponds to a dimer of dimer, the crosslinking events being more favorable in the dimer due to the proximity of BS3 reacting groups.

### The dynamics of P1 and P2 oligomers

To determine if P1 is a condensate of P2 assemblies, dilution experiments were performed prior to solubilization and SEC analysis. As typically shown for the 263K, LA21K *fast* and LA19K strains, P1 oligomers progressively disassembled into smaller oligomers eluting at the elution volume corresponding to the P2 peak position (Fig. 4A-C). For these three strains, the P1 dissociation started with a dilution factor of two, and this dissociation was concerted with a progressive translation of the P1 peak toward the P2 peak (Fig. 4A-C). Similar results were obtained with 22L, Fukuoka-1, ME7 and 139A prion strains (Fig. 4D), indicating that oligomeric assemblies in P1 result from the polymerization of smaller oligomers eluting at the P2 peak position. The fact that P2 from the seven prion strains tested have the same elution volume or partition coefficient (K_AV_) (Fig. 4E) indicates an equivalence for P2 oligomer hydrodynamic size and thus consequently an identical quaternary structure.

**Figure 4:**
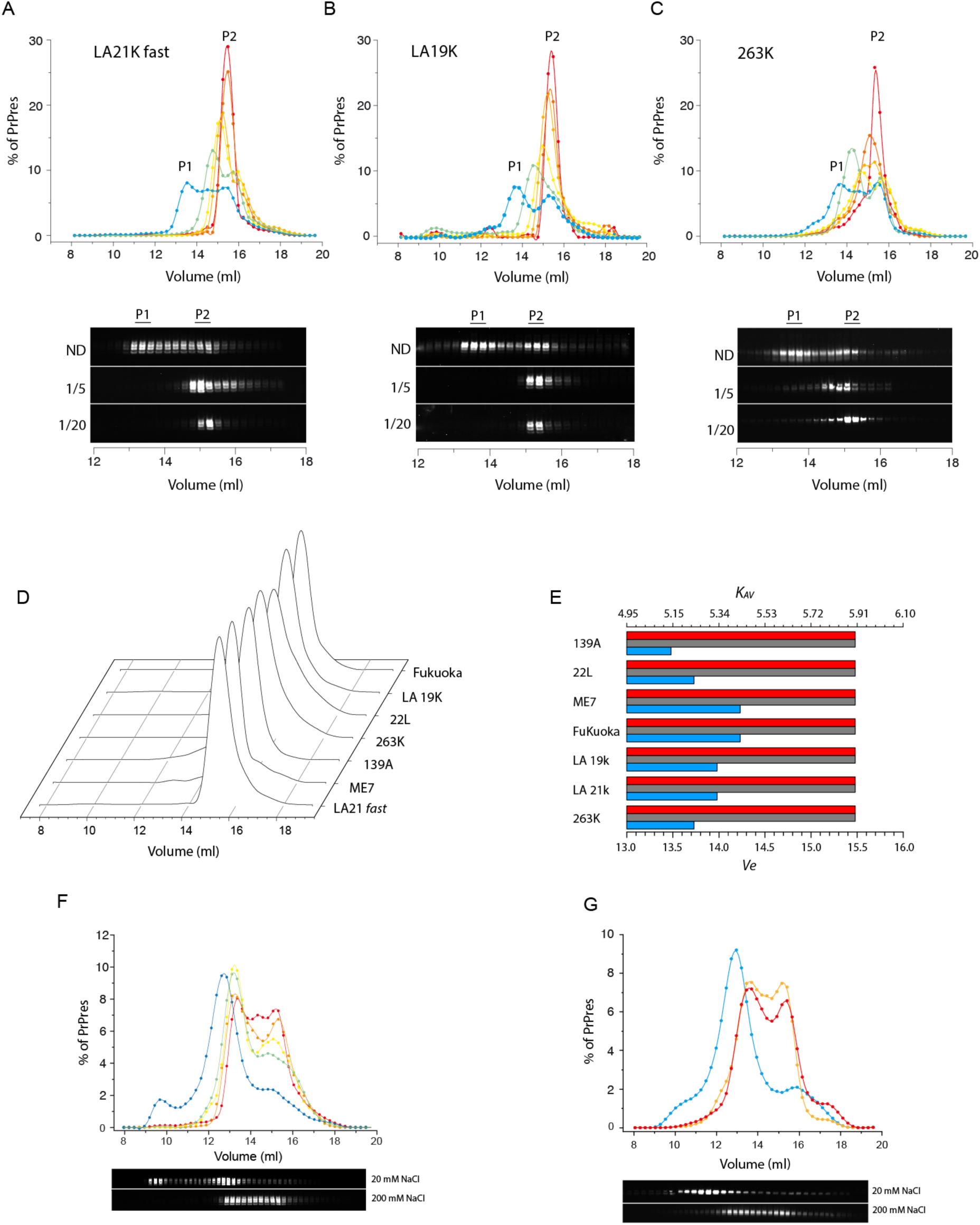
Effect of dilutions on the quaternary structure of P1 and P2 assemblies. (A-C) SEC chromatograms showing the effect of dilution on the size distribution of PrP^res^ assemblies from LA21K *fast*, LA19K and 263K strains. From blue to red, the dilution factors are 1, 2, 3, 5, 10 and 20. Representative western blots used for PrP^res^ level quantification are presented below the chromatograms. (D) Chromatograms representing the effect of a 10-fold dilution factor for seven different prion strains. (E) Representation of chromatogram peak positions reported as the volume of elution (*Ve*) or average distribution constant (*K_AV_*) for the seven prion strains. The gray and blue bar graphs represent the P2 and P1 peak positions before dilution, respectively (Fig. 2A-C), while the red bar graphs represent the peak positions after 10-fold dilution. (F-G) Dependency of the quaternary structure of PrP^res^ assemblies from LA21K *fast* (E) and 263K (F) on the ionic strength of the solubilization media. The blue curve corresponds to ionic strength relative to 20 mM NaCl, and the red curve corresponds to 200 mM NaCl. Representative western blots used for PrP^res^ level quantification are presented below the chromatograms.

The depolymerization of P1 into P2 induced by dilution tends to indicate that weak interactions are involved in the cohesion of the P1 quaternary structure. To increase the strength of these interactions and to investigate the possibility of P1 and P2 condensing into larger assemblies, we explored the effect of ionic strength variations (from 200 mM to 20 mM NaCl) on the quaternary structure of P1 and P2. As shown for 263K and LA21K *fast*, the decrease in ionic strength caused the disappearance of P2, a shift in the size of P1 and the formation of very large assemblies (size >400 kDa). These observations indicate that i) P2 oligomers can polymerize into larger assemblies, and ii) as variations in ionic strength substantially affect PrP size distribution, weak electrostatic interactions govern the condensation dynamics.

### Specific infectivity and replicative propensity of P1 and P2 oligomers

To determine the specific infectivity of the PrP^Sc^ assemblies populating P1 and P2, pools of SEC fractions corresponding to the P1 and P2 peaks from 263K, LA21K *fast* and LA19K prions were intracerebrally inoculated into adequate reporter transgenic mice. By considering the short incubation times in reporter mice for 263K and LA21K *fast* [16], two dilutions were inoculated. The Kaplan‒Meier curves describing the survival percentage as a function of time post-inoculation are depicted in Fig. 5A. P1 and P2 from the three strains induced disease at the full attack rate. However, for the three strains and the two dilutions tested for 263K and LA21K *fast*, the P2 incubation time was significantly shorter than the P1 incubation time. For these three strains, the estimation of the specific infectivity of P1 and P2 based on the ad hoc dose response curves [16] revealed that the specific infectivity of P2 assemblies was 15-100-fold higher than that of P1 assemblies (Fig. 5B). On the other hand, the strain phenotypes of P1 and P2, as assessed by their PrP^res^ electrophoretic signature (Fig. 5C) and regional distribution of PrP^res^ in the brain (Fig. 5D), were globally superimposable and resembled those of the parental strains [21, 31]. The difference in specific infectivity indicates that the structural properties of the assemblies present in the P1 peak are different from those present in P2 to confer distinct replicative properties. The structural difference between P1 and P2 also indicates that P1 does not result from a simple condensation of P2 but a transformation that requires a structural rearrangement.

**Figure 5:**
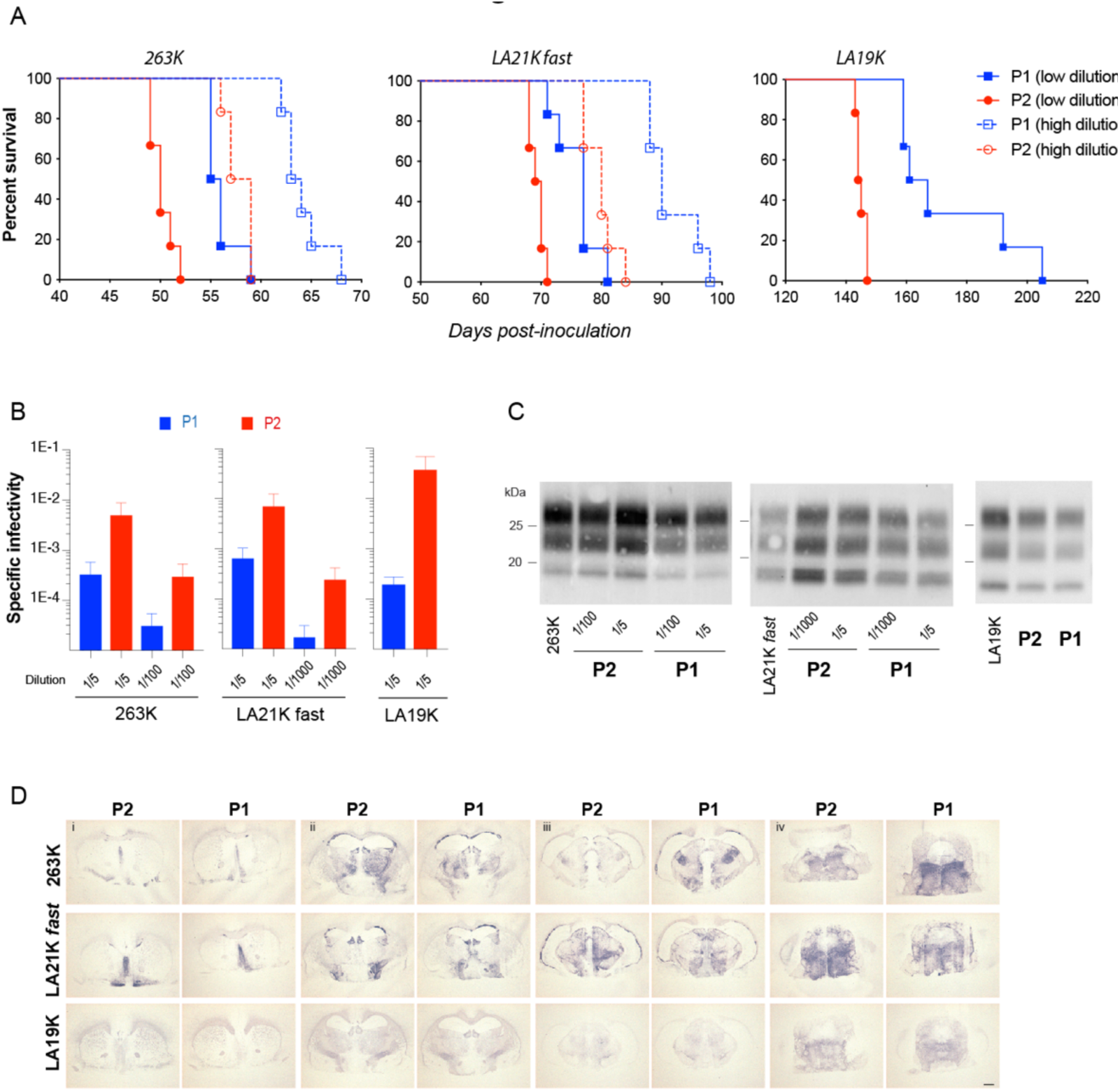
Biological activity of the P1 and P2 assemblies. P1 and P2 assemblies from 263K, LA21K *fast* and LA19K prions were intracerebrally inoculated into reporter tg7 mice expressing hamster PrP (263K) and tg338 mice expressing ovine PrP (LA21K *fast*, LA19K). (A) Kaplan‒Meier curves plot the percentage of mice without prion disease (survival) against the incubation time (days postinoculation). The blue and red colors correspond to the inoculation of P1 and P2, respectively. For 263K and LA21K *fast,* P1 and P2 were inoculated at low and high dilutions (plain lines correspond to 1:5 dilution for all three strains; dashed lines correspond to 1:100 dilution for 263K or 1:1000 dilution for LA21K *fast*). The difference between the P1 and P2 survival curves was statistically significant according to the Mantel‒Cox test at all dilutions tested. (B) Specific infectivity of the P1 and P2 peaks post-SEC fractionation, as calculated from the mean survival time of mice using dose‒response curves. The differences in the specific infectivity values were statistically significant (p<0.05, Mann‒Whitney test). (C) Electrophoretic pattern of PrP^res^ in the brains of mice inoculated with P1 and P2 assemblies. The pattern found after inoculation of unfractionated material is shown for comparison. (D) Representative histoblots of rostro-caudal transversal brain sections after challenge with P1 and P2 assemblies from 263K, LA21K fast (1/5 dilutions) and LA19K prions. Analyses were performed at the level of the septum (i), hippocampus (ii), midbrain (iii) and brainstem (iv). Histoblots were probed with 3F4 (263K) and 12F10 (LA21K *fast*, LA19K) anti-PrP monoclonal antibodies, as indicated. Scale bar, 1 mm.

### Urea unfolding reveals the existence of two sets of assemblies

To further ascertain the structural differences between P1 and P2, we explored the evolution of the P1 and P2 peaks as a function of the concentration of a chaotropic agent such as urea. As shown for LA21K *fast*, LA19K, and 263K, a global quaternary structure rearrangement was observed as a function of urea concentration (Fig. 6A-C). Increasing urea concentration led to a progressive disappearance of P2 and P1 oligomers. Unexpectedly, this was concerted with the formation of a set of higher molecular weight assemblies called P0^U^ with an elution volume lower than P1 and a set of assemblies called P1^U^ with an elution volume slightly higher than P1. The four other prion strains behaved similarly to urea treatment even if the size and amount of P0^U^ was strain dependent, suggesting a common structural rearrangement dynamic (Fig. 6D). Such differential segregation of PrP^Sc^ assemblies during urea exposure is a hallmark of the existence of two sets of assemblies behaving differently.

**Figure 6:**
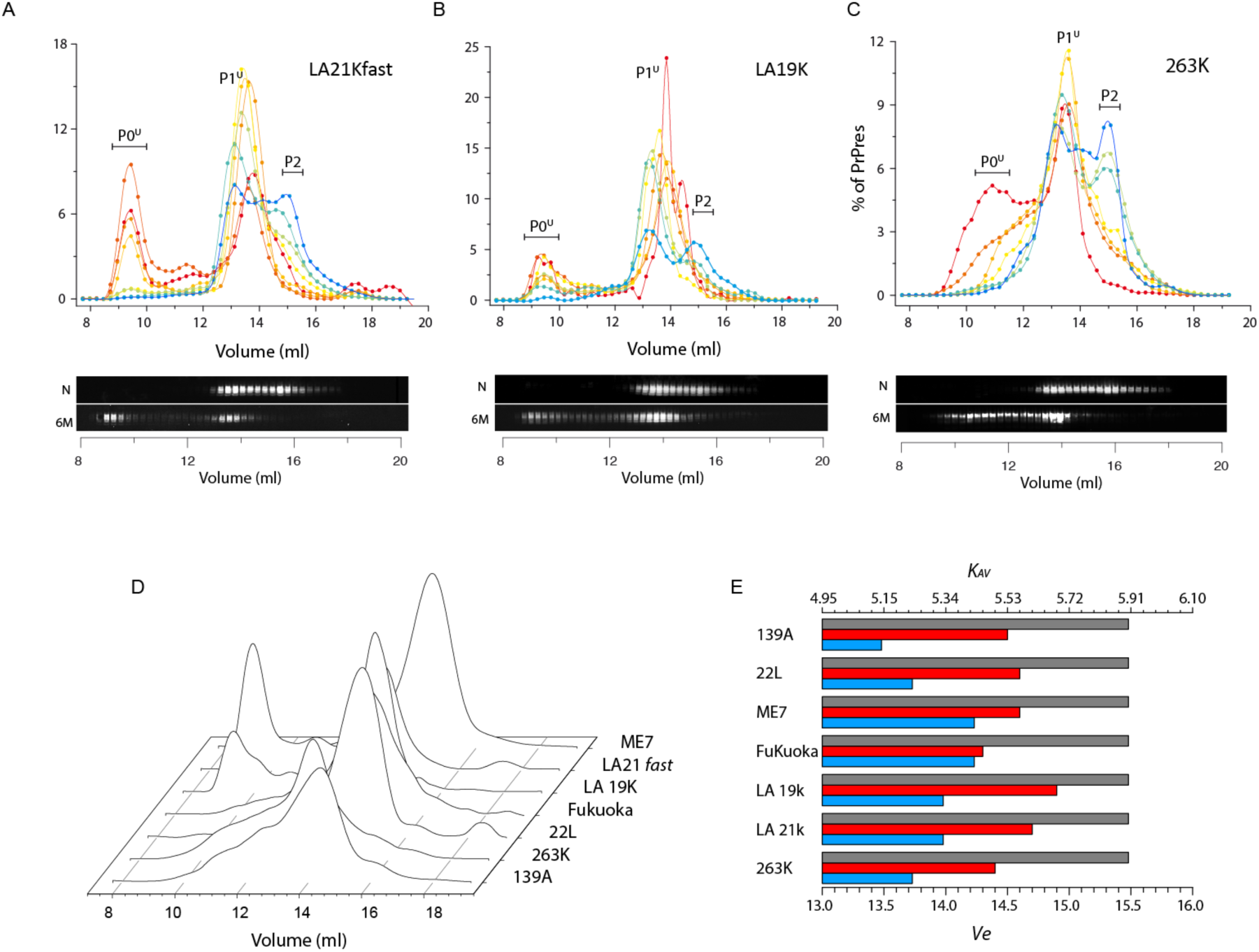
Effect of partial unfolding on the quaternary structure of PrPres assemblies. Typical chromatograms representing the effect of increasing urea concentration on the size distribution of PrPres assemblies for LA21k fast (A), LA19K (B) and 263K (C) strains. For the three strains, the P2 peak progressively disappears as a function of increasing urea concentration in favor of the emergence of a new peak called P0^U^ corresponding to very large assemblies eluting almost at the void volume of the column and P1^U^ eluting at the same elution volume as P1. For each strain, typical western blots of SEC fractions corresponding to native conditions (N) and 6 M urea are presented. Chromatograms corresponding to the effect of 6 M urea treatment on four more strains are also presented (D). For the previous three strains, 6 M urea treatment of 139A, 22L, Fukuoka and ME7 was conducive to the formation of larger assemblies P0^U^ and P1^U^. E) Representation of chromatogram peak positions reported as the volume of elution (*Ve*) or average distribution constant (*K_AV_*) for the seven prion strains. The blue and red bar graphs represent the P1 and P1^U^ peak positions, respectively (Figure 6A-D). For comparison, the peak position of P2 is also represented (in gray).

Among the prion strains tested, the P0^U^ and P1^U^ of LA21K *fast* are the best separated by SEC and are in quasiequal amounts at 6 M urea (Fig. 7A). We thus used this strain to further evaluate the templating activity as well as the biochemical differences between P0^U^ and P1^U^. The infectivity and templating activity were estimated by bioassay in tg338 reporter mice and by PMCA using the tg338 mouse brain as a substrate (Fig. 7B and C). As shown, P1^U^ induced disease at a 100% attack rate, albeit with a delayed mean incubation time of 104 ±2 days, compared to P1 and P2. Reporting this value to the LA21K *fast* dose‒response curve allowed us to calculate that P1^U^ peak infectivity was equivalent to that present in 1.4×10^5^ diluted LA21K *fast* brain material. It also indicated that P1^U^ was >200-fold and >2500-fold less infectious than P1 and P2, respectively. Mice inoculated with P0^U^ did not develop the disease up to 250 days postinoculation (Fig. 7B). The *fast* incubation time of LA21K at the limiting dilution dose (10^-7^) is approximately 150 days [21], indicating the absence of infectivity in P0^U^ as measured by this bioassay. Further analysis of P0^U^ templating activity by PMCA, which is a more sensitive method [32], showed that P0^U^ seeding activity was at least 1000-fold lower than that of P1^U^ (Fig. 7C).

**Figure 7:**
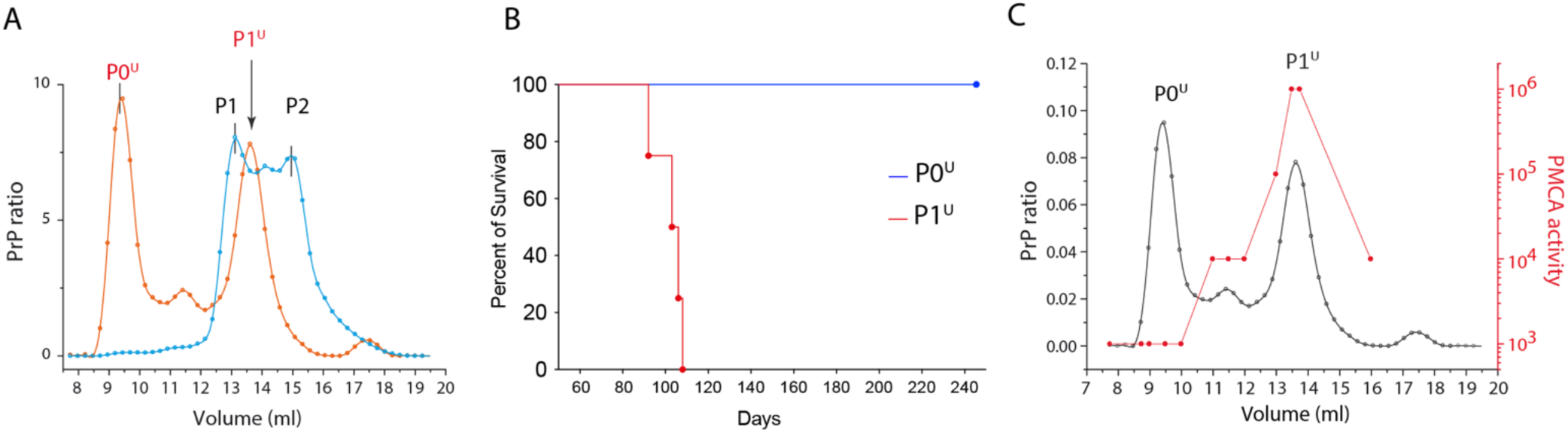
Specific infectivity of P0^U^ and P1^U^ from LA21K fast. Chromatogram showing the separation between P0^U^ and P1^U^ for LA21k fast at 6 M urea (in red). Comparison chromatogram indicating the P1 and P2 positions under native conditions (in blue). (B) Survival of sheep tg338 mice intracerebrally injected with fractions corresponding to the P0^U^ peak (in blue) and P1^U^ peak (in red).

Both the bioassay and the PMCA assay indicate that P0^U^ and P1^U^ have different infectivity and replicative activity and thus are structurally different. These findings further confirm the existence of two structurally distinct sets of assemblies initially present within LA21K *fast* that evolved differently during urea treatment.

## Discussion

The structural diversity of prion assemblies can be qualitatively considered at two scales. At the interindividual scale, it defines the prion strain or prion field isolate. The extent of this diversity at least equals the number of prion strains identified thus far [8, 33, 34]. At the strain scale, the structural diversity reflects the coexistence of PrP^Sc^ subpopulations (or substrain conformers) [23, 26]. The actual dogma considers the strain information encoded in the structure of PrP^Sc^ assemblies when intrastrain structural variations can be defined as the structural diversity of PrP^Sc^ assemblies within a strain. How and in which PrP protein domain(s) such broad diversity could be structurally encoded in a stable manner during multiple replication events remains an entirely open question. The propagation of almost 20 different prion strains in tg338 mice expressing sheep PrP reported as an example in Fig. 1-well demonstrates this paradox.

### A commune dimeric quaternary structure harbors the strain structural determinant

We previously reported that partial unfolding of PrP^Sc^ assemblies is conducive to the formation of a small oligomeric object called suPrP harboring replicative and strain information [35]. In the present work, we show that in biochemical conditions that can be considered native in terms of infectivity and the fold of strain structural determinant [16, 18, 26–28], PrP^Sc^ assemblies spontaneously dissociated into two sets of small infectious oligomeric species, which are called P1 and P2 with respect to their respective elution volumes by SEC. The determination of the molecular weight of the assemblies in the P2 peak by three distinct methods (static light scattering, covalent crosslinking, and hydrodynamic radius estimation) tend to suggest a dimeric quaternary structure. Bioassays in relevant transgenic mice demonstrated that P1 and P2 species conserve replicability and strain properties.

Currently, there is a consensus that both prion infectivity and SSD are encoded in the PrP^Sc^ amyloid fibril fold, even if other quaternary arrangements are increasingly evocated to harbor replicative propensity [31, 35, 3 6]. The fact that small dimeric species such as P1 and P2 harbor both infectivity and strain structural determinants shows that prion replicative properties are not restricted to the fibrillar fold and can also be contained in nonfibrillar arrangements of PrP. We demonstrate that the disassembly process and the dimeric structure of P2 are conserved for seven different prion strains from three distinct mammalian PrP species. Based on the diversity of the tested strains, the different host PrP sequences and the reproducibility of the results, we consider that the disassembly process and the P2 dimeric quaternary structure are generic for all mammalian prions. Such unicity inevitably allows us to conclude that the strain information is not defined by the size of the PrP^Sc^ elementary brick as we previously hypothesized [35] but is defined by the conformation of this dimer. However, considering the growing number of prion strains (considering natural and experimental strains roughly close to 50 [8, 25, 33, 34]), it is not trivial to imagine how such diversity could be encoded in a commune dimeric quaternary structure in a stable manner during multiple replication events without affecting the stability of the oligomerization interface. Such a thermodynamic paradox can be circumvented if the oligomerization domain (i.e., dimerization interface) is structurally independent from the domain encoding the strain information. According to this hypothesis, the domain harboring the SSD could be qualified as a variable domain, differing from one strain to another, while the oligomerization domain remains structurally quasi-invariant from one strain to another. The hypothesis of the existence of two independent folding domains finds a solid structural basis in the comparison of the cryo-EM structures of 263K, aRML, a22L and ME7 (Fig. 8A and B) [9]. Based on these structures, the PrP 95-168 residues adopt a quasicommune fold for the four strains, while the fold of the segment corresponding to 176-231 varies from one strain to another (Fig. 8B). The folding independence of the 95-168 and 169-231 domains is also evidenced in how they contribute to the stabilization energy of the asymmetric unit. Typically, in the case of aRML strain, the total energy of the asymmetric unit is quasiequal to the sum of regions 95-168 and 169-231 (see SI 3). This linear combination highlights the quasiabsence of intradomain interactions (except entropic contributions) through which both domains could influence each other [37, 38]. The quasi-independence of the polymerization domain from SSD also constitutes the best explanation in the limit of current strain typing methods of why different PrP^Sc^ subpopulations within a given strain, such as fibrillar and nonfibrillar PrP^Sc^ [26, 36, 39], harbor the same strain information despite their quaternary structure difference.

**Figure 8:**
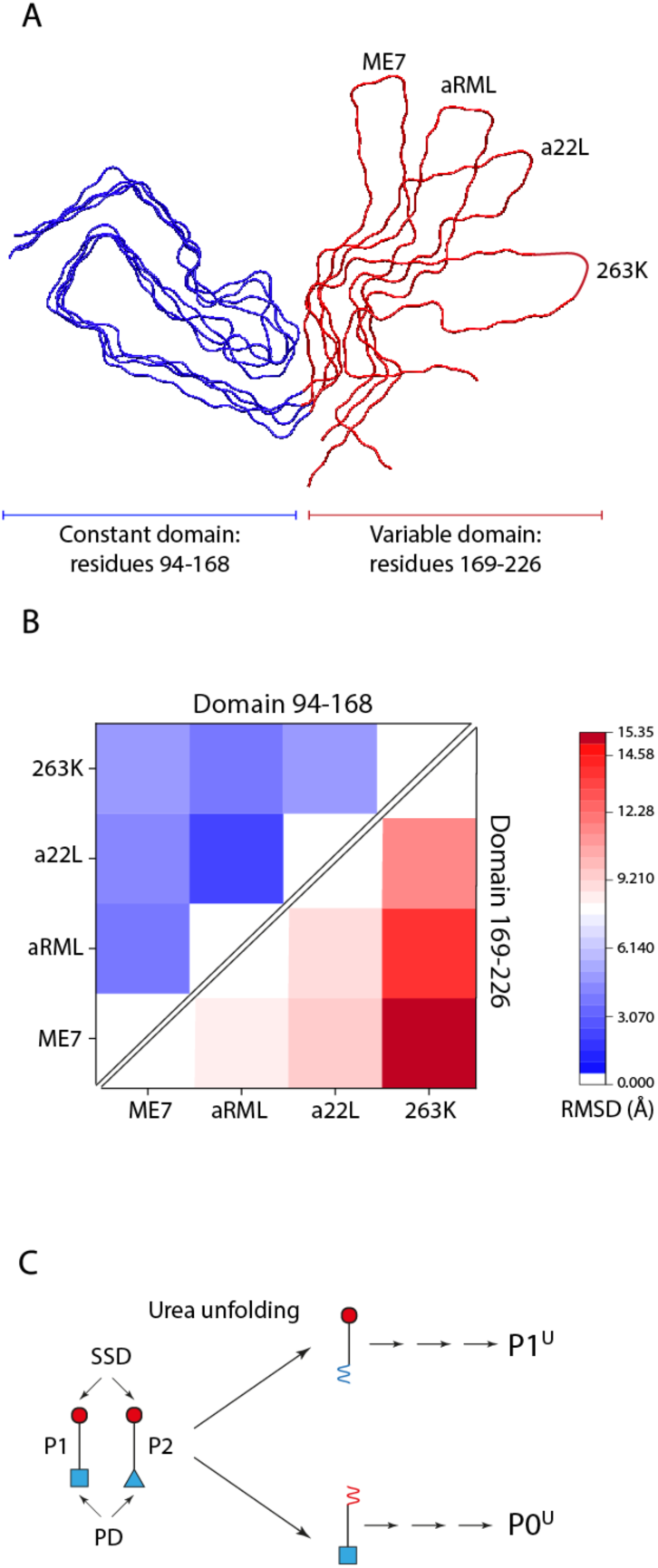
(A) Cryo-EM structures of ME7, aRML, a22L and 263K prion strains represented in an isometric perspective, showing that the domain corresponding to PrP segment 94-168 is quasi-equivalent between the strains while the domain corresponding to the 169-226 region differentiates each strain. These two regions can be respectively qualified as constant domain and variable domain. (B) The RMSD matrix of ME7, aRML, a22L and 263K confirms the quasi-invariance of the 94-168 region and emphasizes the high variability of the 169-226 domain. (C) Hypothetical mechanism of how urea unfolding induces the evolution of P1 and P2 toward P0^U^ and P1^U^. Urea treatment induces a different structural rearrangement in PrP^Sc^ assemblies’ polymerization domain (PD), leading to the formation of two new quaternary structures P0^U^ and P1^U^. The formation of two distinct set of assemblies P0^U^ and P1^U^ with highly significant differences in there biochemical and biological properties can be explained if one considers that the SSD domain of P2 is more sensitive to urea unfolding leading to P0^U^ while the SSD of P1 is more stable and resists to urea unfolding, leading to P1^U^.

### Existence of two conformationally distinct sets of assemblies

Bioassays with relevant transgenic mice and *in vitro* templating activity measurements systematically showed that the specific infectivity and the templating propensity of P1 assemblies from 263K, LA21K *fast* and LA19K prions differ substantially from P2, even if the general strain properties are conserved. These differences are the hallmark of the existence of at least two structurally distinct PrP^Sc^ subpopulations within prion strains. Urea-induced unfolding experiments confirmed this structural polydispersity. The process of urea unfolding gives rise to the formation of two quaternary distinct sets of PrP^Sc^ assemblies. The progressive disappearance of P2, a shift in P1 toward P1^U^ and the formation of large assemblies P0^U^ indicate a profonde rearrangement of the quaternary structure of PrP^Sc^ assemblies upon urea treatment. Even if P1^U^ and P0^U^ amounts and their respective Kav are strain specific, the seven prion strains analyzed here behave similarly during urea unfolding, suggesting that the existence of two intrastrain subpopulations is a generic property of mammalian prions.

In the case of LA21K *fast* prions, where P1^U^ and P0^U^ are well separated by SEC, the measures of infectivity and templating activity indicate that P1^U^ is infectious, while P0^U^ is not infectious and has very low PMCA-templating activity. Only the initial existence of two structurally distinct sets of assemblies could give rise to the formation of two quaternary distinct sets of assemblies, P1^U^ and P0^U^, with vast differences in infectivity, templating activity, and quaternary structure. To explain the specific biochemical properties of P0^U^ and P1^U,^ the P1 and P2 structural domains involved in quaternary structure organization and SSD should have different stabilities with regard to urea denaturation (Fig. 8C). Since the formation of P0^U^ correlates with the disappearance of P2, one can hypothesize that during urea unfolding, the SSD of P2 assemblies could unfold, releasing structural constraints that will allow the modification of the quaternary structure and the formation of larger assemblies as P0^U^. This specific unfolding of the SSD causes the resulting assemblies to lose their strain characteristics and infectivity. In contrast, the formation of P1^U^ could result from the partial unfolding of the polymerization domain of P1 due to its lower stability. This partial unfolding is accompanied by a structural rearrangement of the PD domain of P1, leading to a more compact oligomer P1^U^. This change in compactness also explains the slight shift in the elution volume (or K_AV_) of P1^U^ compared to P1, as systematically observed for all strains tested here.

The existence of two structurally distinct PrP^Sc^ within a given prion raises the question of how they can be generated during the process of prion replication and how they can be maintained. We previously reported that the early step of replication is associated with deterministic structural diversification, giving rise to two specific subpopulations called PrP^Sc^A and PrP^Sc^B [26, 31]. During the evolution of the pathology, PrP^Sc^A disappears in favor of PrP^Sc^B according to a secondary templating pathway that requires the presence of PrP^C^. Even if proof of the equivalence between these two early generated species and P1 and P2 is not established, we show here that whatever the prion strains tested, the structural diversity continues to be maintained throughout the evolution of the pathology, indicating a coevolution of two structurally distinct subpopulations. Bioassay experiments demonstrated that P1 assemblies present lower specific infectivity and templating activity than P2 assemblies, suggesting that during multiple replication events, P2 species should be selected based on the best replicator selection principle. The only way (and nonother exist) to escape the best replicator selection principle is the existence of a pathway of spontaneous or assisted transformation of P2 to P1.

### The high dynamicity of PrP^Sc^ assembly contrasts with a canonical amyloid structure

The effects of dilution and ionic strength on the quaternary structure of P0, P1 and P2 highlight the dynamicity of PrP^Sc^ assemblies. This dynamic balance between a highly aggregated state and the dimeric state fundamentally contrasts with the canonical amyloid fold of prion infectious particles. The cryo-EM structures reveal a mono-brin amyloid organization with protomers in a parallel-in-register intermolecular β-sheet (PIRIBS)-based fold [9–12]. Small oligomeric species associated with or in the vicinity of 263K fibrils were also observed on cryo-EM images [11], highlighting the coexistence of at least two species. As we show here, purified 263K PrP^Sc^ assemblies spontaneously evolve (thermodynamically speaking) from a highly aggregated state into dimeric P2 species. Even if the condition of this transformation requires the presence of detergents, such as dodecyl maltoside, sarkosyl and incubation at 37 °C, it highlights the existence of a transformation pathway between PrP^Sc^ amyloid fibrils and the dimeric P2 species that does not significantly affect the original SSD but deeply affects the quaternary structure organization.

Two hypotheses can explain the relationship between amyloid fold and dimeric P2 species. The first hypothesis is that amyloid assemblies result from the simple condensation of the dimeric objects P2. An alternative hypothesis would be a spontaneous transformation of amyloid fibril assemblies into dimeric P2 species by a structural rearrangement of fibrillar PrP^Sc^. As P2 species harbor the infectivity and all the attributes of the strain, the transformation should not affect the SSD but only the domain involved in the organization of the quaternary structure. The availability of a high-resolution structure for 263K amyloid fibrils can help to distinguish between these two hypotheses. According to the cryo-EM structure of RML PrP assemblies at 2.70 Å resolution [11], the estimation of the standard free energy of the trimeric asymmetric unit of the monofilament using the AMBER force field (see SI3) is approximatively −14000 kcal/mol. For comparison, the standard free energy of sheep recombinant PrP is approximately −9 kcal/mol [40], and a carbon‒carbon covalent bond and a disulfide bridge are approximately −60 kcal/mol and −58 kcal/mol, respectively [41]. It clearly appears that in comparison to the unfolding energy of recombinant PrP or to a covalent C-C and S‒S bond, the stability of the RML PrPres monofilament is high enough to render these assemblies unresponsive to a simple equilibrium displacement by dilution or change in ionic strength at 37 °C. The high stability of the PIRIBS-based fold argues in favor of a spontaneous transformation of amyloid fibrils into P2 species and discards the condensation hypothesis. The even-more-complex-to-tackle transformation hypothesis may explain why highly purified 263K amyloid filaments spontaneously generated P2 without any P1, as reported in Figure 3.

## Conclusion

The high dynamicity of PrPSc assemblies highlighted in the present work contrasts with the deadpan dogma of PrPSc assemblies. We demonstrated that amyloid fibrils are not the unique quaternary structure organization harboring prion infective determinants, but alternative PrP^Sc^ assemblies such as small oligomeric species such as P1 and P2 also harbor infectivity and SSD. This duality led us to propose a folding-domain separation between the strain determinant and folding domain involved in the formation of the quaternary structure. The existence of such an alternative infectious quaternary structure also questions the principle of templating and fibril end-elongation as a unique mode of prion replication.

## Materials and Methods

### Ethics

Animal experiments were conducted in strict accordance with ECC and EU directives 86/009 and 2010/63 and were approved by the local ethics committee of the author’s institution (Comethea; permit numbers 12/034, 15/045 and APAFIS#29603-2021020914525215).

### Brain homogenate preparation and solubilization

Stocks of infected brain homogenate (BH, 20% w/v in 5% glucose from Syrian hamsters for the 263K strain, from tg338 transgenic mice for the LA21K *fast* and LA19K strains, and from tga20 transgenic mice for the 22L, Fukuoka-1, ME7 and 139A strains diluted to were treated with 80 µg/mL PK for 1.5 hrs at 37 °C under gentle agitation. The reaction was stopped by the addition of Pefabloc at a final concentration of 2 mM in the BH. Prior to size exclusion chromatography (SEC), 200 μl BH was mixed with an equal volume of 200 μl solubilization buffer (2X concentrated) to reach a final concentration of 25 mM HEPES (stock 0.5 M, pH 7.4), 150 mM NaCl, 10 mM EDTA, 5 mM n-dodecyl β-D-maltoside (DM) and 50 mM sarkosyl. The mixture was then incubated at 37 °C for 4 hrs under gentle orbital agitation (300 rpm on Eppendorf Thermomixer) and centrifuged at 15 000 × g for 5 min prior to SEC analysis. Urea unfolding experiments were performed by providing 200 μl of BH followed by the addition of urea (solid) to the solubilization buffer to reach 1 to 7 M final urea concentration in the BH after the addition of solubilization buffer to the BH. As the addition of urea, specifically at high concentrations, causes a change in the sample volume, the amount of added solubilization buffer was adjusted to always ensure a constant concentration of the components of the solubilization in the final sample.

### Size exclusion chromatography and quaternary structure determination

SEC analysis was performed using an ÄKTA-100 purifier FPLC (series 900, Amersham Biosiences, Amersham, UK) and a Superdex 200 10/300 GL gel filtration column (24 ml, GE, Chicago, US). The composition of the running buffer was 25 mM HEPES and 150 mM NaCl, and the pH of the buffer was adjusted to pH 7.2. For ionic strength and urea experiments, the NaCl and urea concentrations of the running buffer and solubilized BH were identical. In all SEC experiments, no detergents were added to the running buffer to avoid the formation and maintenance of micellar structures [28, 29]. After centrifugation at 10 000 × g for 3 min (no visible pellet), the entire sample solution was loaded on an SEC column using a sample loop. During elution, fractions of 250 µl were collected at 0.35 ml/min. Between each SEC run, the column was sanitized with three times the column volume of a 2 M sodium hydroxide. The column was calibrated using blue dextran molecules with varying molecular weights between 10 and 300 kDa, standard globular proteins (Bio-Rad), PrP^C^ and recPrP. The mean average molecular weight (*<MW>*) as well as size distribution prior to any treatment of purified 263K [30] assemblies was estimated by a homemade static light scattering (SLS) device and by dynamic light scattering (DLS, Malvern). The <MW> of fractions obtained from SEC analysis of purified 263K after solubilization was determined according to the Rayleigh relation by normalizing the SLS intensity by the ratio of PrP [42, 43]. Crosslinking was performed by incubating fractions corresponding to the P1 and P2 peaks with the 11 Å bifunctional amine crosslinker bis-sulfosuccinimidyl suberate (BS3) at final concentrations of 0.5 mM and 2 mM for 15 min at 37 °C. The reaction was stopped by the addition of 1 mM Tris-buffer and analyzed by conventional western blot using Sha31b antibody.

### Dilution experiments

For each prion strain, BH (20% w/v in 5% glucose) was first treated with 80 µg/mL PK for 1.5 hr at 37 °C under gentle agitation. The reaction was stopped by the addition of Pefabloc to a final concentration of 2 mM. Then, the PK-treated BH was diluted by a given factor (1/2, 1/3, 1/5 and 1/10) in 5% glucose solution v/v prior to adding 2X solubilization buffer and incubation at 37 °C for 4 hrs under gentle orbital agitation (500 rpm on Eppendorf Thermomixer). The product was then analyzed by SEC as described earlier.

### PrP quantification by western blot

Western blotting was used to quantify the amount of PrP in SEC fractions. Samples were deposited onto 26-well 12% Bis-Tris Criteroin XT precasted gels (Bio-Rad Laboratories Inc., Hercules, CA, USA) and electrotransferred onto a nitrocellulose membrane with a semidry electrotransfer system (Schleicher & Schuell BioScience, Whatman). Immunoactivity was probed with biotinylated Sha31 antibodies [44] for 15-20 min at room temperature. For all SDS‒PAGE analyses, a fixed quantity of human PrP was employed for consistent calibration of the PrP signals in different gels. To improve the sensitivity of the western detection method for samples containing low levels of PrPres, double deposition was performed to electroconcentrate the sample. Typically, after a first round of sample loading in SDS‒PAGE wells, a short 2 min migration was performed to allow running within the acrylamide gel for 2 mm followed by a second round of sample loading before the migration was continued until the front reached 3 cm within the gel. The electrotransfer and detection remained unchanged.

### Miniaturized bead-PMCA assay

Protein misfolding cyclic amplification was conducted according to a method first developed by Soto et al. [45] and further optimized in-house by Moudjou et al. [46]. In brief, the PrP^C^ substrate for incubation with PrP^Sc^ was obtained from Tg7 mice [16] overexpressing hamster PrPC. Up to 12-month-old mice were euthanized with CO_2_, and brains were recovered and washed with Ca^2+/^Mg^2+^-free PBS containing 5 mM EDTA. Next, the PrP^C^ lysate (10% (w/v)) was prepared in PMCA buffer (50 mM Tris-HCl at pH 7.4, 5 mM EDTA, 300 mM NaCl, 1% Triton X-100) by Dounce cooling in ice with 20-30 strokes to ensure complete homogenization of the brains. Up to 9 brains were mixed to avoid biased results due to anomalies in individual mice. The lysate was incubated for 30 min at 4 °C before centrifugation at 1000 g at 4 °C for 2 min. The supernatant was frozen at −80 °C until further use. PMCA detection was performed in a 96-well plate using a final volume of 40 µl. Four microliters of the sample was mixed with 36 µl of Tg7 lysate, and one Teflon beaker (diameter 2.38 mm, Marteau et Lemarié, Pantin, France) was added. From this solution, seven serial 10-fold dilutions were generated. The final concentration depended on the initial concentration of the sample in the lysate. Starting with an already diluted sample enables the application of higher maximal dilution factors. The microtiter plate was closed and placed in an S300 sonicator (Delta Labo, Colom- belles, France) and subjected to 96 cycles of 30 s sonification (30% amplitude) followed by 30 min of incubation at 35 °C. PMCA requires the rigorous use of controls to avoid false-positive results. Therefore, a tg7 lysate was used as a negative control, and an already positively tested 263K lysate was used as a positive control. The generation of PrP^Sc^ during PMCA was verified by the presence of PK-resistant PrP. Hence, products of the PMCA run were supplemented with SDS (0.3%) and treated with PK (115 µg/ml final concentration) at 37 °C for 1 hr. The digestions were stopped by the addition of Laemmli denaturation buffer 2x and subsequent heating at 100 °C for 5 min. The content of PrPres was determined by dot blot analysis. In a 96-well plate, 20 µl of the PK-treated PMCA product was mixed with 50 µl of SDS solution (1%) and glycerol to a final glycerol concentration of 20% (w/v). The samples were transferred to a nitrocellulose membrane using a dot blot apparatus, and immunoactivity quantification was performed according to the same procedure described for western blot analysis.

### Bioassays

The pool of fractions of interest was extemporarily diluted tenfold in 5% glucose and immediately inoculated via the intracerebral route into reporter mice tg338 and tg7 (20 μl per pool of fraction, n = 5 mice per pool). Mice showing prion-specific neurological symptoms were euthanized at the end stage. To confirm the presence of prion disease, brains were removed and analyzed for PrP^Sc^ content using the Bio-Rad TsSeE detection kit prior to immunoblotting, as described above. The survival time was defined as the number of days from inoculation to euthanasia. To estimate the influence of the difference in the mean survival times, strain-specific curves correlating the relative infectious dose to survival times were used, as previously described.

## Acknowledgments

This work was supported by grants from the Fondation pour la Recherche Médicale (Equipe FRM DEQ20150331689), the European Research Council (ERC Starting Grant SKIPPERAD, number 306321), the Ile de France region (DIM MALINF) and the support of Hochschule Fresenius GEM. GMBH university.

---

**S1.**
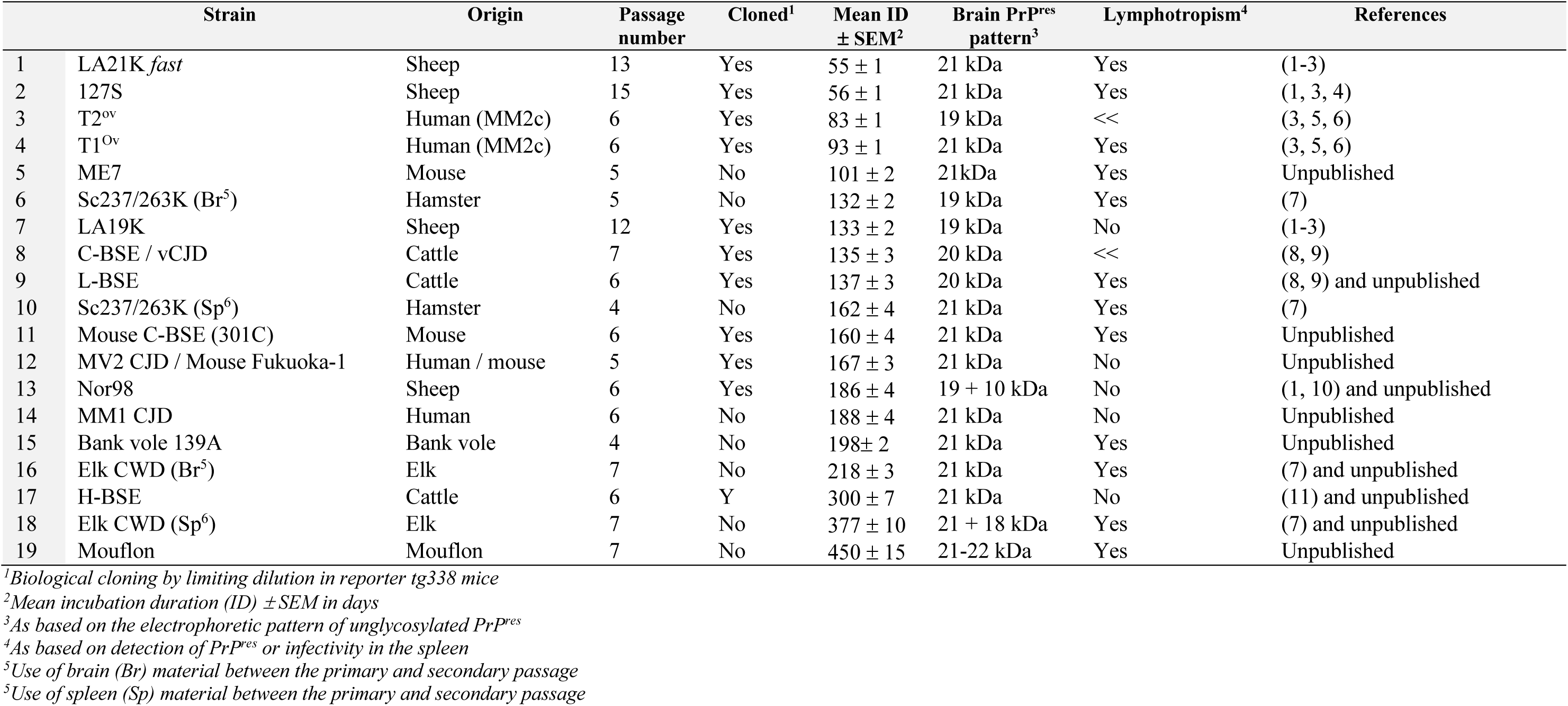
List of stable prion strains with differing biological or biochemical phenotype upon adaptation to tg338 mice.

**S2:**
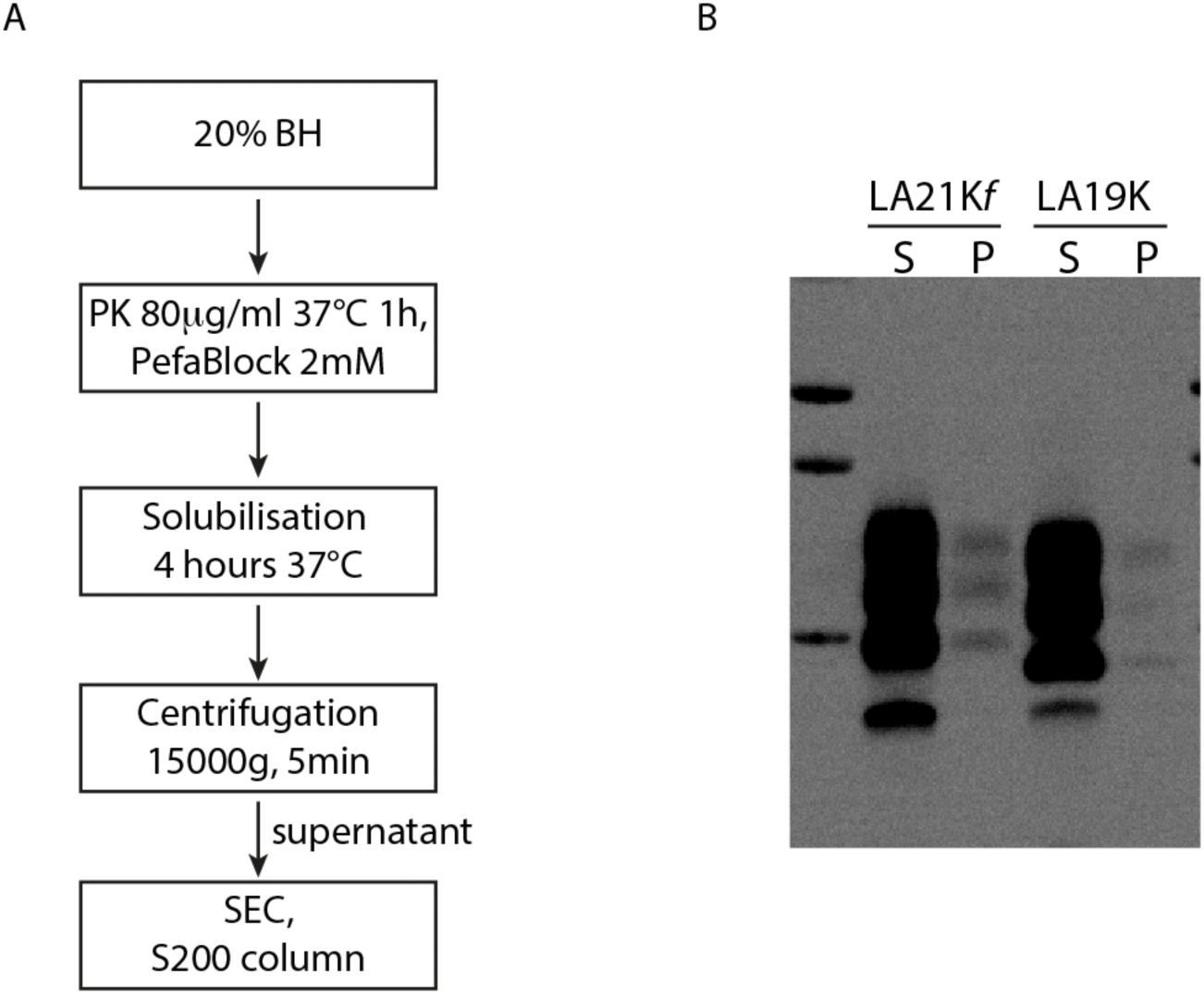
Procedure of brain homogenate solubilization. (A) The solubilization procedure of the brain homogenate prior SEC. As described in the material and method 20% BH of the end-stage infected mice is first treated by PK to remove the reminiscent PrP^C^. The PK was then inactivated by PefaBlock and diluted two-fold in the solubilization buffer and incubated for four hours at 37°C under gentle agitation. At this stage the solubilized BH has a translucid appearance. A five minutes centrifugation at 15 000g leads only to the formation of submillimeter pellet. The PrP amount estimation in the supernatant versus pellet has performed by western blot by resuspending a pellet obtained for the solubilization of 200ul of brain homogenate 20% directly in 20ul of 2X-Laemeli buffer for western blot analysis. The supernatant was 2X diluted in 2X-Laemli buffer. As typically shown for LA21Kfast and LA19K strain the amount of PrP in the pellet is neglectable compared to the supernatant. This observation indicates that most of the PrPSc assemblies have been solubilized by this procedure.

**S3:**
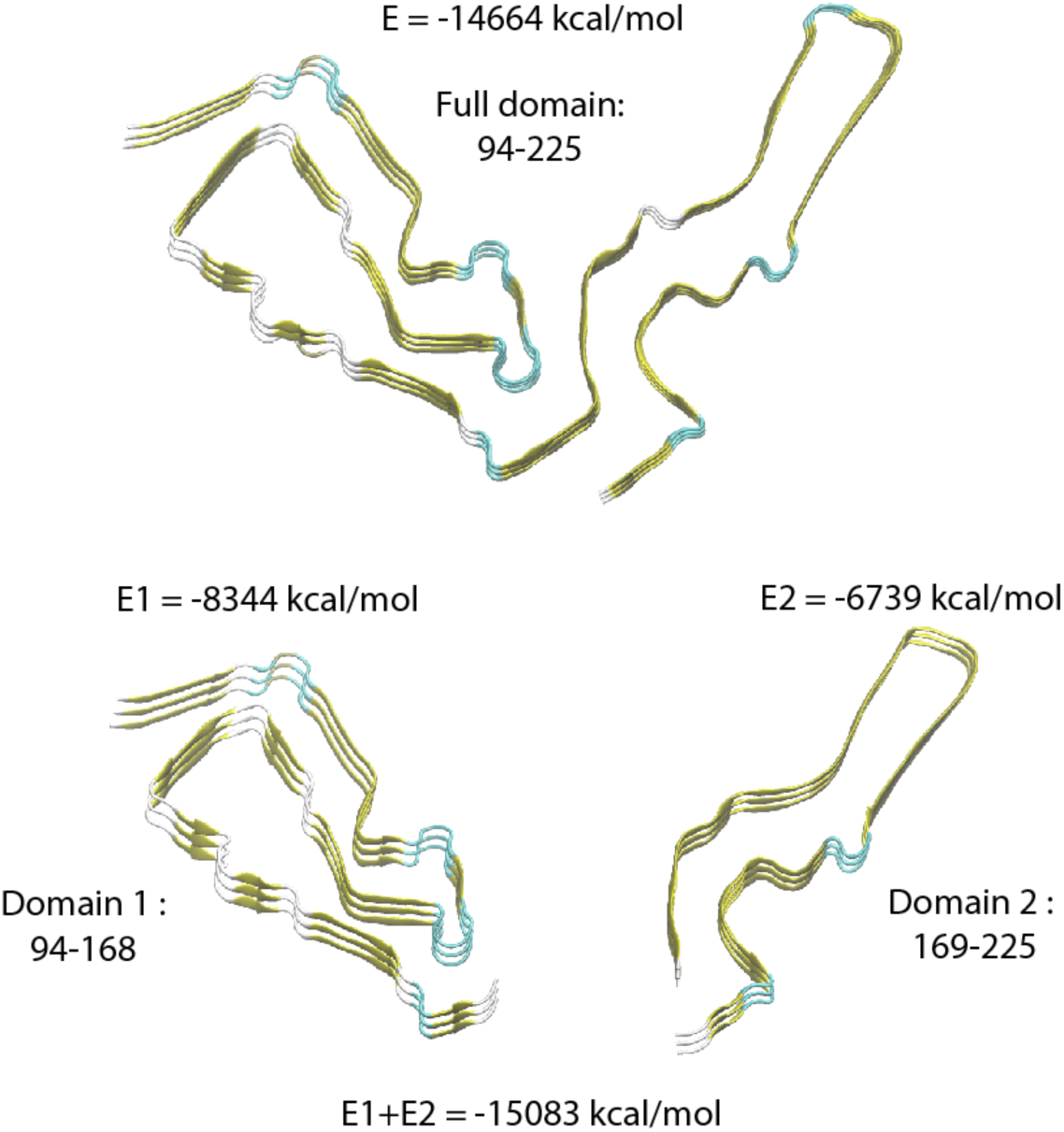
Energetic of the asymmetric unit of RML prion strain PrPres. We explored the stabilization energy of the asymmetric unit of RML PrPres prion strain (PDB: 7quig) as well as its domain 1 (residues 94-168) and domain 2 (residues 169-225). We first proceed to their structural refinement by preforming molecular dynamics in the presence of implicit water using an AMBER force filed ff14Sb. The energy calculation (see the above figure) of the full domain as well as the isolated domain 1 and 2 revealed that the total energy of the asymmetric unit is quasi-equal to the sum of the energy of the region 95-168 and 169-231. This linear combination highlights the quasi-absence of intra-domain interactions through which both domains could influence each other. The folding independency of these two domains has also for consequence that in certain specific physico-chemical conditions one domain could undergo into a rearrangement without affecting the other domain.

## Notes

### Competing Interest Statement

The authors have declared no competing interest.

